# Sox11 overexpression restores embryonic pro-growth transcription in mature corticospinal tract neurons

**DOI:** 10.1101/2025.09.30.679327

**Authors:** Elizabeth Batsel, Zimei Wang, Elizabeth Otten, Rabia Mohammad, Paula Pascual, Darby O’Shea, Jose Rosas, Pantelis Tsoulfas, Murray G. Blackmore

## Abstract

Neurons in the central nervous system (CNS) display a high capacity for axon growth during early development but lose this ability at a pivotal differentiation stage marked by synaptic maturation, circuit integration, and a profound shift in gene transcription. Once mature, most CNS neurons fail to reverse this transcriptional switch after axon injury, fundamentally constraining their intrinsic capacity for axon regeneration. Here, we show with single-nucleus RNA sequencing that forced expression of the transcription factor Sox11 in mature corticospinal tract (CST) neurons produces large-scale and stable changes in gene expression that are highly enriched for growth-relevant processes, and which strongly resemble those of pre-synaptic embryonic stages. Moreover, Sox11 is equally effective when delivered to chronically injured CST neurons. These data reveal the ability of Sox11 to reverse a critical step of neuronal maturation even in otherwise unperturbed neurons, clarifying the transcriptional underpinnings and highlighting the potential of Sox11 to act as a pro-regenerative stimulus.

## INTRODUCTION

A recurring observation in developmental and regenerative biology is that as cells progress through maturation, they often lose the capacity for regeneration. This tradeoff is particularly striking in the mammalian central nervous system (CNS), where embryonic and neonatal neurons display a high capacity for axon regeneration that is lost during early development [1–4]. Neuronal maturation is a staged process including neurogenesis, migration, axon elongation, axon and dendritic arborization, and prolonged synapse refinement [4]. Of these, the transition from axon elongation to arborization and synapse formation is of particular interest, as it appears to coincide with the pronounced reduction in the capacity for long-distance regenerative axon growth [4,5]. Transcriptional comparisons of neurons in the elongation stage to more mature counterparts have revealed pronounced differences including strong upregulation of transcripts involved in ion transport and synaptic vesicle cycling and downregulation of transcripts involved in cytoskeletal regulation, cell adhesion, and axonal transport [6–10]. These transcriptional changes are likely major drivers of the maturational loss of regenerative axon growth ability, making their reversal an important objective in pro-regenerative research.

Multiple lines of evidence suggest the transcription factor Sox11 is a central regulator of axon growth ability. Developmentally, Sox11 is widely expressed in CNS and peripheral nervous system (PNS) neurons during the stage of axon elongation, is sharply downregulated in postnatal periods around the transition to arborization/synaptogenesis, and has been shown in loss-of-function experiments to be necessary for embryonic axon elongation [11–14]. Sox11 is also strongly linked to regenerative axon growth. For example, mammalian dorsal root ganglion (mDRG) neurons and zebrafish retinal ganglion cells (zRGCs), which display a high capacity for spontaneous axon regeneration, strongly upregulate Sox11 in response to axon damage [15–19]. Mammalian retinal ganglion cells (mRGCs), which display minimal spontaneous regeneration but robust axon elongation in response to molecular interventions such as PTEN knockdown, also spontaneously upregulate Sox11 after injury [20,21]. Loss of Sox11 interferes with mDRG peripheral regeneration and almost completely blocks mRGC regeneration stimulated by PTEN knockdown, demonstrating a critical contribution of Sox11 re-expression to the regenerative process [22,23]. Conversely, forced elevation of Sox11 in mDRG and mRGC neurons after axon damage triggers upregulation of regeneration-associated genes such as *doublecortin* (Dcx) and downregulation of transcripts involved in synapse function, leading to enhanced regenerative axon growth [24,25]

Although these experiments establish a function for Sox11 in neurons that are actively engaged in a broader injury response, it is unknown whether Sox11 possesses the ability to induce expression of regeneration-associated genes and/or trigger embryonic reversion under stable, homeostatic conditions. This question is of critical importance for regenerative efforts that focus on neuronal subtypes which normally display smaller responses to axon damage and for neurons in a chronic injury state after the initial injury response has subsided. For example, we recently showed that corticospinal tract (CST) neurons, an important sensorimotor population with low regenerative capacity, undergo only minor transcriptional changes after spinal injury, likely owing to the large distance between injury and cell body [26]. This raises the question of how transcriptional stimuli like elevated Sox11 might affect transcription in a less injury-perturbed context.

Here we employed single-nucleus sequencing to determine the transcriptional consequences of forced expression of Sox11 in CST neurons. We show in both uninjured and chronically injured CST neurons that Sox11 triggers a wide-spread transcriptional response marked by upregulation of cytoskeletal-interacting proteins previously linked to axon growth and down-regulation of transcripts involved in synapse function. In addition, Sox11-sensitive transcripts displayed high overlap with transcripts up- or down-regulated during CST maturation, indicating a partial embryonic reversion. Interestingly, comparison of transcripts affected by Sox11 in adult CST neurons with prior studies in mRGCs and epidermal cells revealed a common gene network with cytoskeletal functions, suggesting a conserved function for Sox11 across diverse cell types. Thus, Sox11 expression is sufficient to drive large-scale transcriptional effects and partial embryonic reversion even in neurons not engaged in an active injury response, highlighting the centrality of Sox11 in neuronal maturation and supporting its potential utility as a pro-regenerative stimulus.

## RESULTS

### Effects of Sox11 Overexpression on CST Transcription

We first used single-nucleus sequencing to determine the transcriptional effects of forced expression of Sox11 in adult corticospinal tract (CST) neurons in homeostatic, uninjured conditions. Following previously established procedures, AAV2-Retro expressing Sox11 or titer-matched control was injected into the cervical spinal cord of adult male and female mice along with nuclear localized H2B-mGreenLantern (H2B-mGl) to facilitate subsequent nuclear isolation [26]. To track transcriptional changes through time, Sox11-treated animals were divided into four cohorts that received unique molecular barcodes, and which were sacrificed at one, two, four, or six weeks after injection. Cortical tissue from at least three animals per time point was collected by fluorescent microdissection, frozen, and then pooled for unified purification of CST nuclei by fluorescence activated nuclei sorting (FANS) and single-nucleus library preparation using a 10X Genomics platform (**Supplementary Fig. 1-1**). Expression of Sox11 was confirmed by immunohistochemistry in one randomly selected animal from each time point (**Supplementary Fig. 1-2**). The resulting pooled data were merged with a separate library derived from control-treated CST nuclei. De-multiplexing of the four barcoded time points was performed in Seurat, allowing each to be compared separately to control-treated nuclei and to one another (**Fig. 1a**).

**Figure 1.**
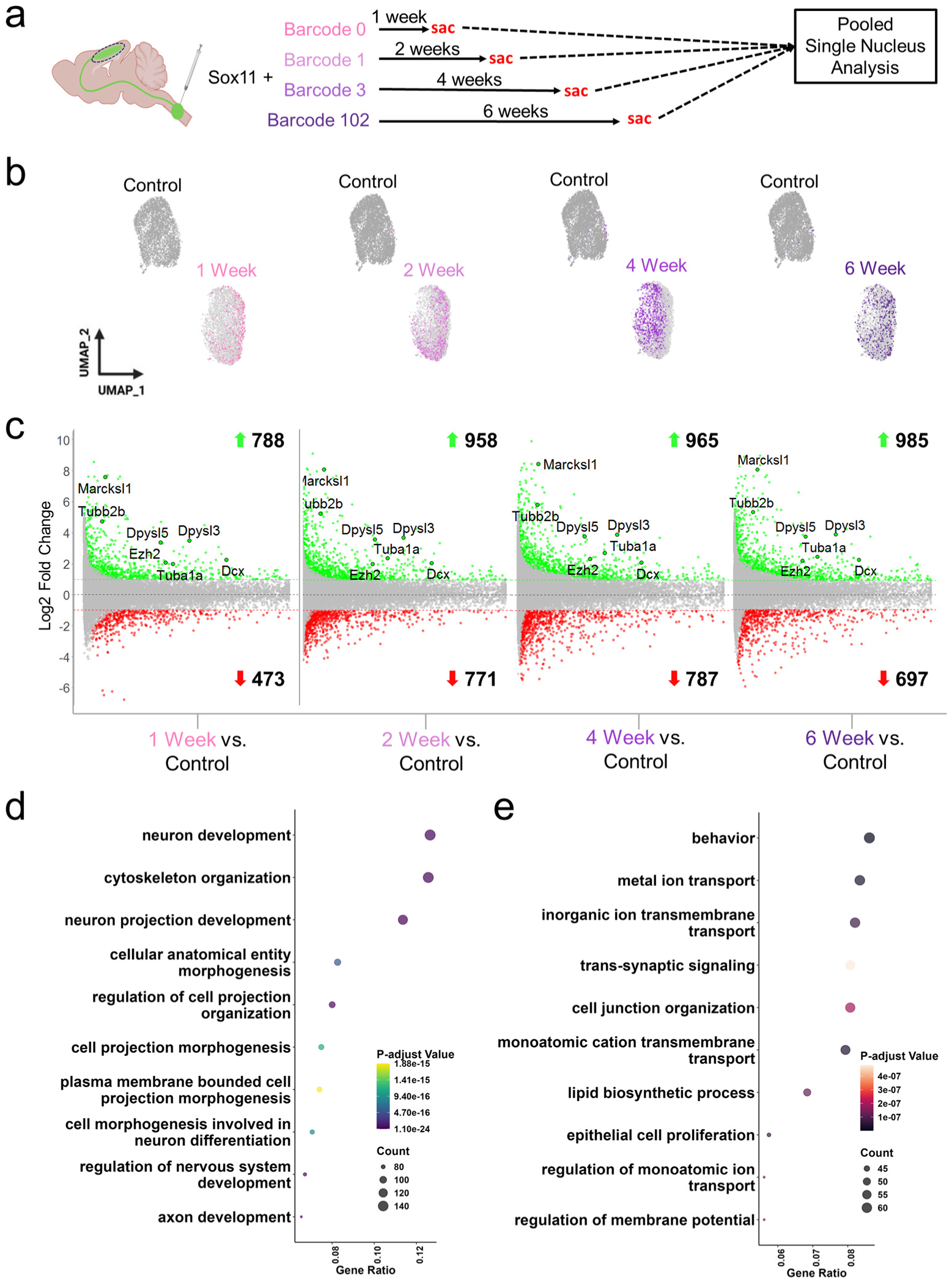
Single-nucleus profiling detects a large transcriptional response to Sox11 overexpression. (a) Experimental design in which CST neurons of adult male and female mice were retrogradely labeled by nuclear-localized AAV2-Retro-H2B-mGreenlantern, AAV2-Retro-Sox11, and a unique molecular barcode. Fluorescently labeled tissue from each group was dissected after one, two, four, or six weeks and pooled for single-nucleus profiling. (b) UMAP visualization shows separation of Sox11-treated nuclei from control and co-clustering of all Sox11 time points. (c) MA plots showing the relationship between transcript abundance and differential gene expression at the four time points, comparing Sox11-treated to control nuclei (non-parametric Wilcoxon rank sum test, p-value < 0.05), which reveal numerous transcripts are significantly up-regulated (green) and down-regulated (red). (d) Gene ontology analysis of transcripts that are significantly upregulated in Sox11-treated nuclei shows enrichment for terms such as neuron development and cytoskeletal organization (e) Gene ontology analysis of transcripts that are significantly downregulated in Sox11-treated nuclei shows enrichment for terms including ion transport and synaptic signaling (p-values indicated in legend, Benjamini-Hochberg test).

UMAP clustering produced two main clusters, one containing control nuclei and the second containing Sox11-treated nuclei from all four time points (**Fig. 1b**). The separation of Sox11 from control suggests a distinct transcriptional state, while the co-clustering of Sox11 groups suggests relative similarity through time. Indeed, this visual impression was confirmed by differential gene (DEG) analysis. Using a standard threshold of two-fold or greater change (Log2FC > 1, p < .05), Sox11-upregulated transcripts numbered 788 by one week, climbed to 958 by two weeks, and remained at comparable levels at four and six weeks (965 and 985, respectively) (**Fig. 1c, Supplementary Table 1-1**). Downregulated transcripts followed a similar pattern, numbering 473 by one week, 771 by two weeks, and 787 and 697 at four and six weeks, respectively (**Fig. 1c, Supplementary Table 1-1**). Two-way comparisons between Sox11 time points revealed some differences between one week and later time points (maximum 179 DEGs between weeks one and six) but very small differences between two weeks and later time points (maximum 15 DEGs between two and six weeks) (**Supplementary Fig. 1-3, Supplementary Table 1-1**). In summary, forced elevation of Sox11 produces large-scale transcriptional changes in uninjured CST neurons that are largely established and stable by two weeks post-transduction.

We next performed gene ontology (GO) enrichment analysis on transcripts up- and down-regulated by Sox11. Top biological processes enriched in Sox11-upregulated transcripts included neuron development, axon development, and cytoskeleton organization (**Fig. 1d, Supplementary Table 1-2**). Strongly upregulated transcripts included *Dpysl3, Dpysl5, Tubb2b, Tuba1a, Marcksl1, Dcx,* and *Ezh2*, which have been functionally linked to axon growth (**Fig. 1c, Supplementary Table 1-1**) [27–30]. Conversely, Sox11-downregulated showed enrichment for terms related to synapse function and the regulation of membrane potential, processes that have been previously linked to the maturational decline in axon growth ability [5,8] (**Fig. 1e, Supplementary Table 1-2**). For example, the downregulated set included *Slc12a5* (Kcc2), and *Cacna2d2*, which have been linked to both neuronal excitability and to axon regeneration [8,31]. Fluorescent *in situ* hybridization on a second cohort of animals confirmed strong upregulation of *Dpysl3*, *Dpysl5*, and *Sox11* in animals that received AAV2-Retro-Sox11 (**Fig. 2**). Combined, these results show Sox11’s transcriptional effects to include an elevation of transcripts linked to axon extension and a diminishment of synapse-related transcripts.

**Figure 2.**
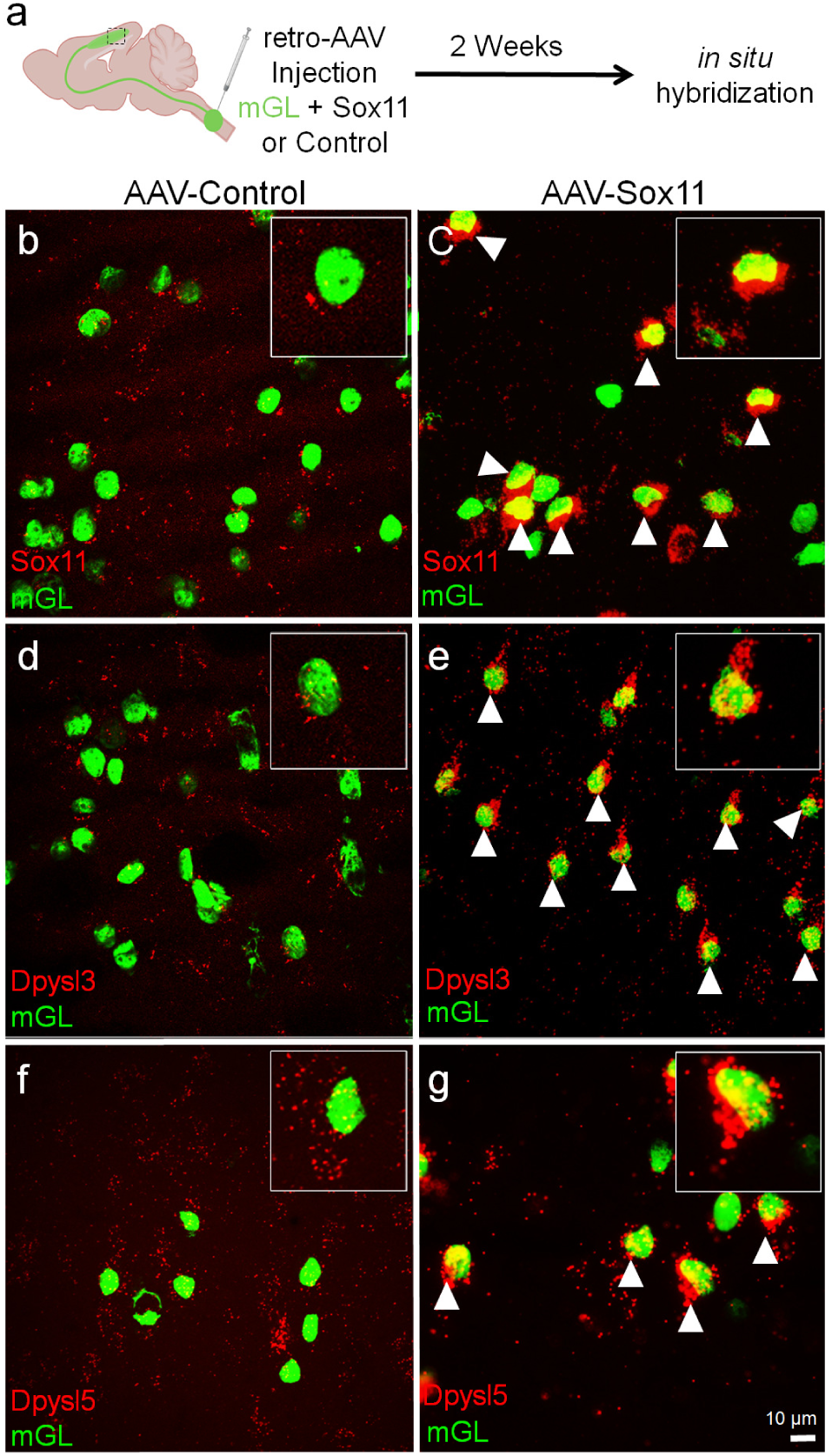
Fluorescence *in situ* hybridization confirms upregulation of Sox11, Dpysl3, and Dpysl5 in Sox11-treated animals. (a) Experimental design involving retrograde transduction of CST neurons with nuclear-localized AAV2-Retro-H2B-mGreenlantern (H2B-mGl) alone or in combination with AAV2-Retro-Sox11, followed 2 weeks later by *in situ* hybridization to detect candidate genes. (b-g) show coronal sections of cortex with CST nuclei labeled by H2B-mGl (green). Sox11, Dplysl3, and Dplysl5 (red) are readily detectable in Sox11-treated animals (c, e, and g respectively) but not in H2B-mGl only animals (b, d, f respectively).

### Sox11’s Effect on Transcription in a Chronic Injury Model

Sox11’s efficacy as a potential pro-regenerative treatment requires an ability to influence transcription in neurons after injury, which could present a unique cellular context. Based on eventual clinical feasibility we focused on the sub-acute to early chronic phase of injury. Adult mice received a contusion injury to the right cervical spinal cord, severing the corticospinal tract. Four weeks after injury, animals received an injection of H2B-mGl and AAV2-Retro-Sox11 or a matched control into the cervical spinal cord just above the injury site. Two weeks later, the left (injured) cortex was harvested, CST nuclei were isolated by FANS, and single-nucleus analyses performed as described above (**Fig. 3a**) [26]. Immunohistochemistry for glial fibrillary acidic protein (GFAP) on spinal sections confirmed interruption of the right dorsal column (**Fig. 3b**). In addition, immunohistochemistry for Sox11 performed on randomly selected treatment and control animals confirmed effective overexpression (**Supp. Fig. 3-1**).

**Figure 3.**
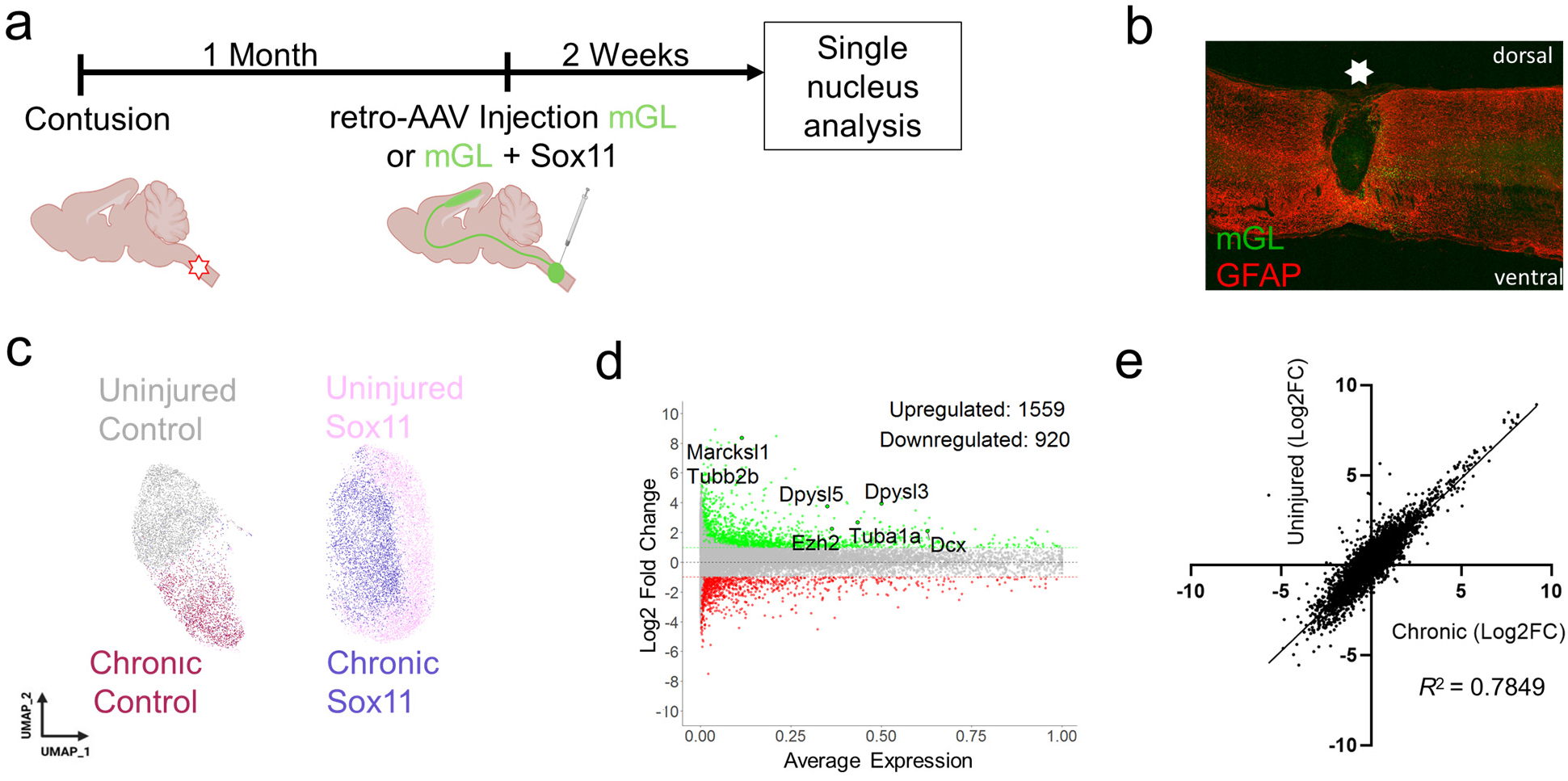
Single-nuclei profiling detects a large transcriptional response to Sox11 overexpression after a chronic contusive spinal injury. (a) Experimental timeline. Adult mice received a unilateral contusion at cervical level five followed one month by retrograde transduction of CST neurons with nuclear-localized AAV2-Retro-mGreenlantern (H2B-mGI) alone or in combination with AAV2-Retro-Sox11. Two weeks later, brains and spinal cords were dissected. (b) Shows a sagittal section of a spinal cord stained for GFAP (red) to confirm interruption of the CST tract in dorsal spinal cord (c) UMAP visualization of Uninjured/Control (grey), Uninjured/Sox11 (pink), Injured/Sox11 (purple), and Injured/Control (maroon) nuclei indicates separation by gene treatment but not by injury state (d) A MA plot of transcripts differentially expressed after Sox11 expression in chronically injured CST neurons shows numerous genes significantly upregulated (green) and downregulated (red) (non-parametric Wilcoxon rank sum test, p-value < 0.05). (e) Linear regression of Log2 fold changes triggered in uninjured versus chronically injured CST neurons reveals strong positive correlation (R-squared = 0.7849, slope 0.99).

The resulting data were merged in Seurat with the prior sets from non-injured CST that received control virus or Sox11 for two weeks, resulting in a unified, four-group object consisting of non-injured/AAV-control, non-injured/AAV-Sox11, injured/AAV-control and injured/AAV-Sox11. UMAP visualization produced two clusters, one containing AAV-control nuclei from both injured and uninjured animals and the other containing AAV-Sox11 nuclei, also of mixed injury status (**Fig. 3c**). Thus, nuclei segregated by gene treatment but not injury status, indicating a dominant Sox11 effect regardless of injury state. Notably, growth-relevant transcripts *Marcksl1, Dcx, Ezh2, Tubb2b, Dpysl3* and *Dpysl5* were upregulated similarly upon Sox11 overexpression in injured CST neurons (**Fig. 3d**, **Supplementary Table 3-1**). Indeed, linear regression analysis of Sox11 DEGs in uninjured versus injured CST neurons showed a striking correlation (R-squared = 0.7849, slope 0.99) (**Fig. 3e**). These results show that Sox11’s ability to drive large-scale transcriptional changes in CST neurons is maintained when delivered four weeks after spinal injury.

### Conserved Activation of Cytoskeletal and Axon Development Networks Across Cell Types

We further contextualized our results by comparing them to prior profiles of other cell types responding to forced expression of Sox11. We first considered a study in murine retinal ganglion cells (mRGCs) that were pre-treated with Sox11 and challenged by optic nerve injury and compared to RGCs that received injury alone [24]. Like the present study, Sox11 was delivered by AAV about two weeks prior to transcriptional profiling, resulting in genome-wide detection of Sox11-responsive DEGs. **Supplementary Table 4-1** reports for each gene symbol the fold change and p-value in mRGCs side-by-side with the corresponding Sox11 effect in CST neurons, facilitating direct comparison. Overall, we found Sox11 to produce a much larger transcriptional response in CST neurons than was previously reported in RGCs. DEGs numbered 4707 in CST vs 1827 in RGC without fold-change threshold and 1699 in CST vs. 766 in RGCs at ±2-fold threshold) (**Supplementary Table 4-1**). To determine whether the identities of Sox11-responsive transcripts were similar in the two cell types we created a set of of Sox11- of responsive transcripts were similar in the two cell types we created a set of high-confidence CST-DEGs with consistent effects across our multiple experiments and then intersected with the prior RGC-DEGs. Downregulated DEGs showed no significant overlap (1.13 enrichment p=0.075, hypergeometric test, **Supplementary Table 4-1**), indicating that Sox11 caused distinct transcripts to decline in each cell type. In contrast, intersection of upregulated DEGs revealed 127 in common, somewhat more than expected by chance (1.52 enrichment, p=6.14×10^-7^, hypergeometric test) (**Fig. 4a, Supplementary Table 4-1**). Gene ontology analysis of these upregulated transcripts showed neuron projection development and axon development among the most enriched terms (**Fig. 4c, Supplementary Table 4-2**), while protein-protein interaction (PPI) analysis revealed a central network highly enriched for cytoskeletal regulation and axon development (**Fig. 4e**). Within a smaller set of 32 transcripts that exceeded two-fold upregulation in both cell types we found *Dcx, L1cam,* and *Stmn2* (SCG10), previously linked to axon growth [28,29,32,33] (**Supplementary Table 4-1)**. Thus, although Sox11’s effects in CST are larger and mostly distinct from those previously reported in RGCs, they include a subset that are commonly upregulated, and which are involved in cytoskeletal regulation and axon growth.

**Figure 4.**
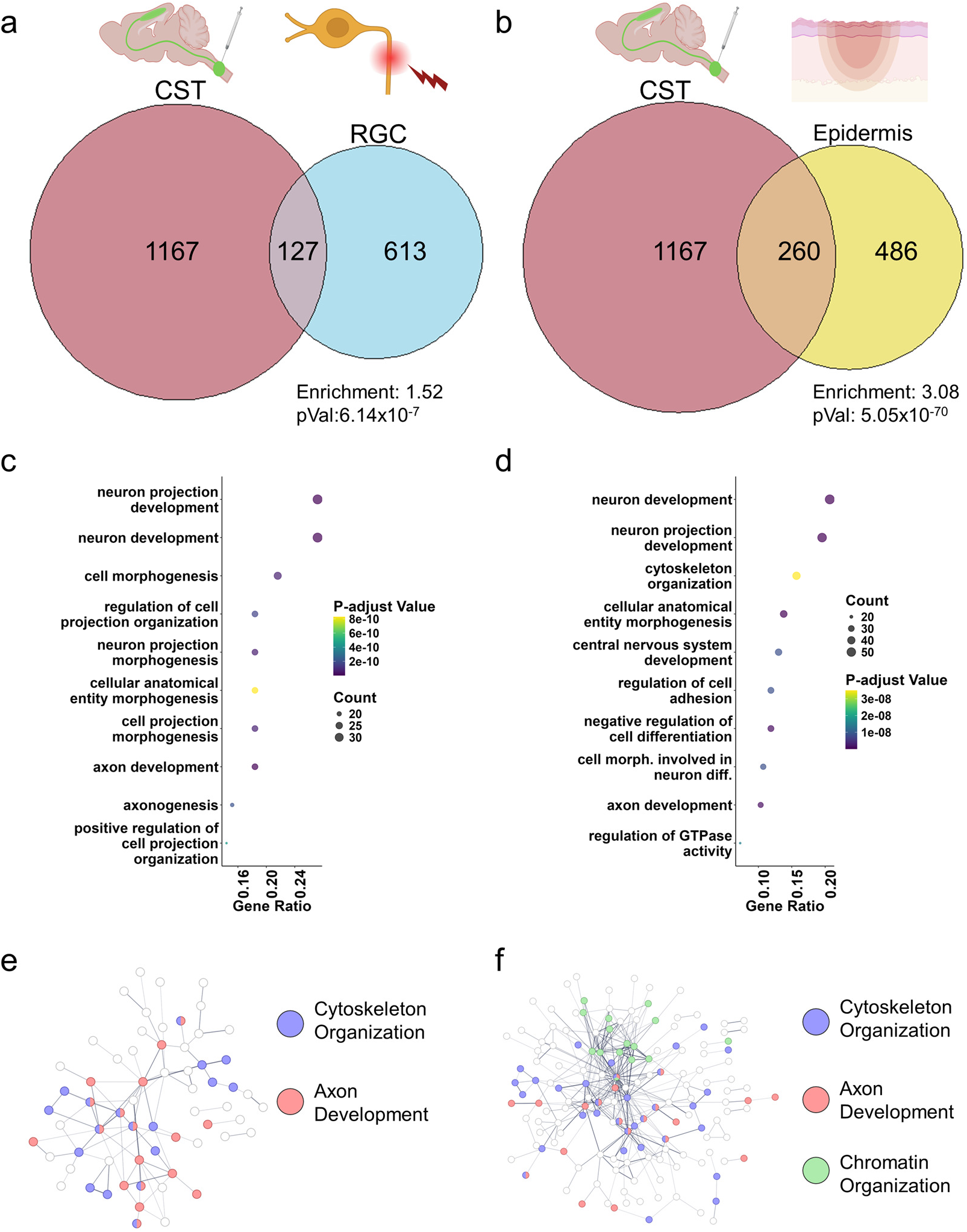
Sox11 activated conserved growth-related networks across cell types. (a) A Venn diagram illustrating the overlap of upregulated genes in CST neurons and retinal ganglion cells (RGCs) following Sox11 overexpression, highlighting both unique and shared transcriptional responses. (b) A Venn diagram shows the overlap of upregulated genes in CST neurons and epidermal cells after Sox11 overexpression, also indicating unique and shared transcriptional responses. P-values were determined by hypergeometric tests. (c–d) Gene Ontology (GO) enrichment analysis for overlapping genes from (a) and (b). (e) An interaction protein network (String) of the genes shared among CST and RGCs after Sox11 overexpression reveals two distinct functional categories: (1) Cytoskeletal Organization and (2) Axon Development. (f) String network of the genes shared among CST and epidermal cells after Sox11 overexpression reveals three distinct functional categories (1) Cytoskeletal Organization, (2) Axon Development, and (3) Chromatin Organization. Each network node represents a gene, with edge represents a known interaction.

We next considered a study conducted in murine epidermal cells, in which forced expression of Sox11 was shown to enhance wound healing and cell motility [34]. Transcriptional profiling found 776 Sox11-responsive transcripts significantly upregulated and 266 downregulated at a 2-fold threshold, which are incorporated into **Supplementary Table 4-1**. Like the mRGC comparison, downregulated transcripts were largely distinct, with only 54 overlapping (p=0..501, hypergeometric test). In contrast, upregulated transcripts showed high similarity between CST and epidermal cells, with 260 transcripts in common, 188 of which exceeded 2-fold upregulation in both datasets (4.22 enrichment, p=2.2×10^-71^, hypergeometric test) (**Fig. 4b, Supplementary Table 4-1**). Reminiscent of the mRGC overlap, GO and PPI analysis showed high enrichment of terms related to cytoskeleton and axon development and a core protein network with similar terms (**Fig. 4d, f, Supplementary Table 4-2**). Commonly upregulated transcripts included various tubulin isoforms, *Marcks, Marcksl1,* and *Dpysl5* (**Supplementary Table 4-1**). This shared transcriptional response across cell types helps validate the current results and indicates that activation of transcripts related to cytoskeleton-based motility is a conserved function of Sox11.

### Sox11 Partially Recapitulates an Embryonic-Like Pattern of Gene Expression

Finally, we probed for similarity between Sox11’s transcriptional effects in CST neurons and transcriptional changes that occur naturally during maturation. The available description of CST maturation is based on microarray technology and includes maximum cell age of only postnatal day 14 [6]. Therefore, to facilitate more direct comparison with our Sox11 profiling, we performed single-nucleus analysis of mouse cortex at embryonic day 18 (E18), a time when cortical projection neurons including CST are actively elongating axons [35]. We prepared two replicate libraries and then used a Seurat-based pipeline to extract nuclei from deep-layer glutamatergic projection neurons from the initially mixed-cortical data. This entailed the identification and removal of off-target cell types including mitotically active progenitors (*Pax6*/*Top2a*), oligodendrocyte precursor cells (*Sox10*/*Pdgfra*), radial glial cells (*Slc1a3*/ *EAAT*), inhibitory neurons (*Gad2*), and shallow layer glutamatergic neurons (*Cux2*) (**Supp. Fig. 5-1**). In the remaining nuclei, *Bcl11b* was used as a positive marker to select deep layer (DL) projection neurons [36]. Finally, the resulting data were merged with the prior adult CST data, visualized by UMAP, and compared to identify DEGs (**Fig. 5a**).

**Figure 5.**
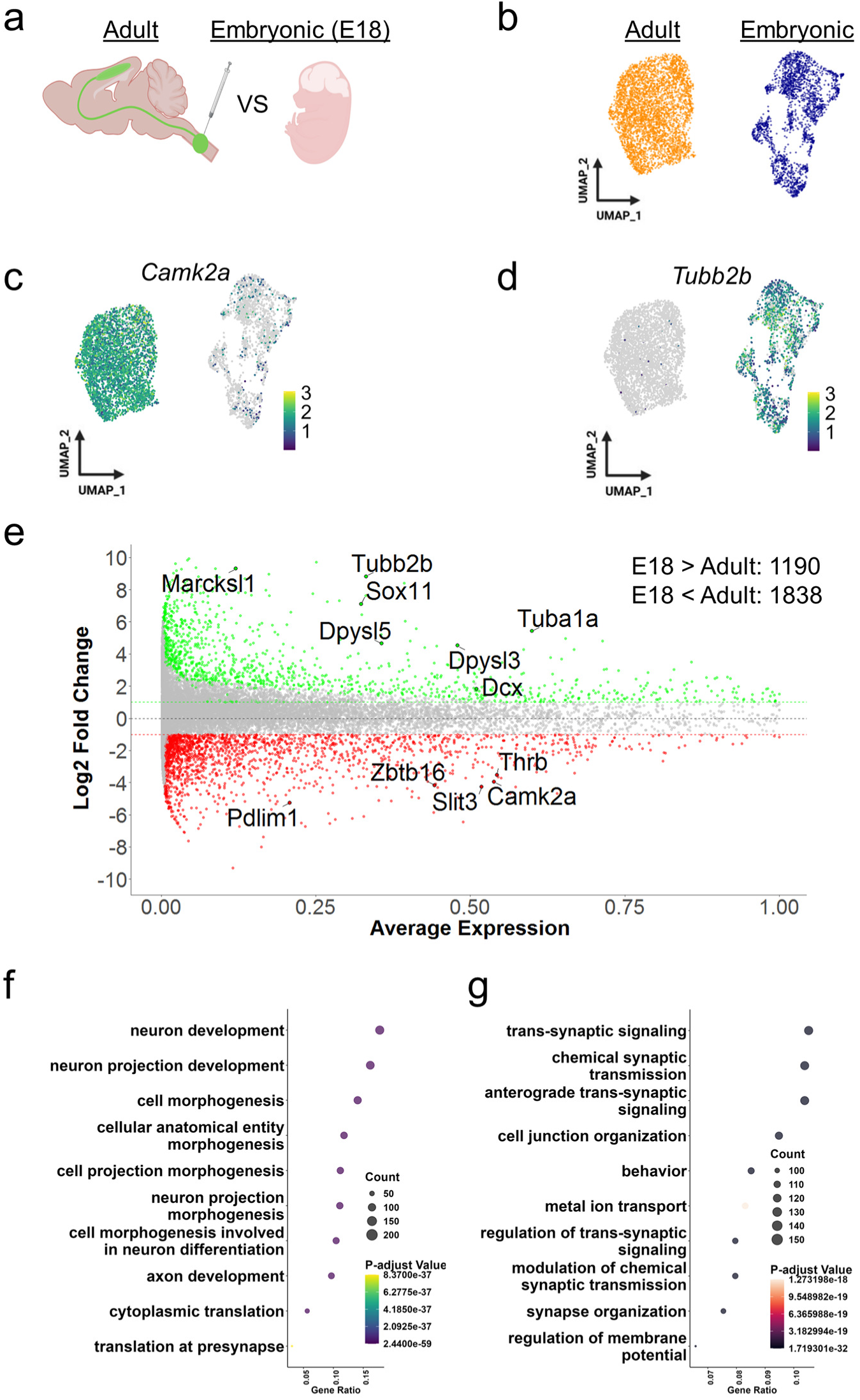
Single-nucleus profiling detects large changes in transcription associated with cortical neuronal maturation. (a) Experimental design showing comparison between adult CST and embryonic cortical tissue. (b) UMAP visualization shows separation of adult (orange) and embryonic (blue) nuclei (c,d) Feature plots of *Camk2a* (c) and *Tubb2b* (d) expression confirm expected enrichment in adult and embryonic nuclei, respectively. in adult versus embryonic nuclei. (e) MA plot showing transcripts significantly more abundant in E18 (green) or in adult CST (red) (2-fold threshold, p-value < 0.05, non-parametric Wilcoxon rank sum test, p-value < 0.05). (f,g) Gene ontology analysis indicating biological processes that are significantly enriched in E18 and adult gene sets, respectively (p-value<0.5, Benjamini-Hochberg test).

Adult CST and embryonic DL nuclei clustered distinctly by UMAP (**Fig. 5b**). For initial validation, we examined transcripts known to be developmentally regulated and observed the expected concentration of *Camk2a* and *Tubb2b* in the adult and embryonic clusters, respectively (**Fig. 5c, d**). DEG analysis identified 1,190 genes more than 2-fold more abundant in embryonic nuclei (hereafter embryonic transcripts) and 1838 that were more abundant in adult (hereafter adult transcripts) (**Fig. 5e, Supplementary Table 5-1**). For additional validation we identified numerous strongly regulated DEGs with concordance to known patterns of developmental up- or down-regulation (indicator transcripts labeled in **Fig. 5e**), and FISH confirmed *Dpysl3*, *Dpysl5*, and *Sox11* to be abundant in E18 cortex but low or absent in the adult (**Supplementary Fig. 5-2**). Gene ontology analysis indicated the enrichment of terms including neuron development and axon development for the embryonic transcripts and ion transport and synaptic signaling for the adult transcripts (**Fig. 5f,g, Supplementary Table 5-2**). Moreover, algorithmic prediction of transcriptional regulation (IPA) yielded Sox11 as a top candidate to regulate the set of embryonic transcripts (Activation Z-score 5.23, p= 4.3×10^-13^, Fisher’s Exact Test) (**Supplementary Fig. 5-3, Supplementary Table 5-3**). These data identify broad transcriptional differences between cortical projection neurons engaged in embryonic axon extension and fully mature CST neurons and provide initial support for Sox11’s involvement in mediating this difference.

We next compared these maturational changes to the previously characterized effects of Sox11 expression in adult CST neurons. Manual inspection revealed many of the previously noted Sox11-responsive transcripts. For example, *Dpysl3, Dpysl5, Dcx, Ezh2, Marcksl1*, and *Tubb2b*, all of which are Sox11-upregulated and linked to axon growth, were also among the most differentially expressed embryonic transcripts (**Fig. 5e, Supplementary Table 5-1**).To test more directly for enrichment of shared genes, we intersected Sox11-upregulated and embryonic DEGs, both selected by 2-fold thresholds and p<.05. This analysis revealed a striking overlap; 475 transcripts were common to both sets, more than four times the number predicted by chance (4.38 enrichment, p=2.14×10^-212^, hypergeometric test) (**Fig. 6a, Supplementary Table 6-1**). Linear regression of this shared set showed a strong positive correlation between fold changes induced by Sox11 and differential expression at E18 (R-squared 0.7199), further supporting a potential role for Sox11 in regulating this subset of embryonic transcripts (**Fig. 6c**). Similarly, intersection of Sox11-downregulated transcripts and adult-DEGs also revealed strong overlap (661, 4.74 enrichment, p=4.61×10^-360^, hypergeometric test) (**Fig. 6b, Supplementary Table 6-1**), although with weaker fold-change correlation (R-squared = 0.1147) (**Fig. 6d**). Finally, GO analysis indicated enrichment of terms related to neuron development and axon development in the shared Sox11-Up/Embryonic set and membrane potential and synaptic transmission in the shared Sox11-Down/Adult set (**Fig. 6e, f; Supplementary Table 6-2)**. Thus, forced expression of Sox11 partially restores in adult CST neurons an embryonic pattern of transcription.

**Figure 6.**
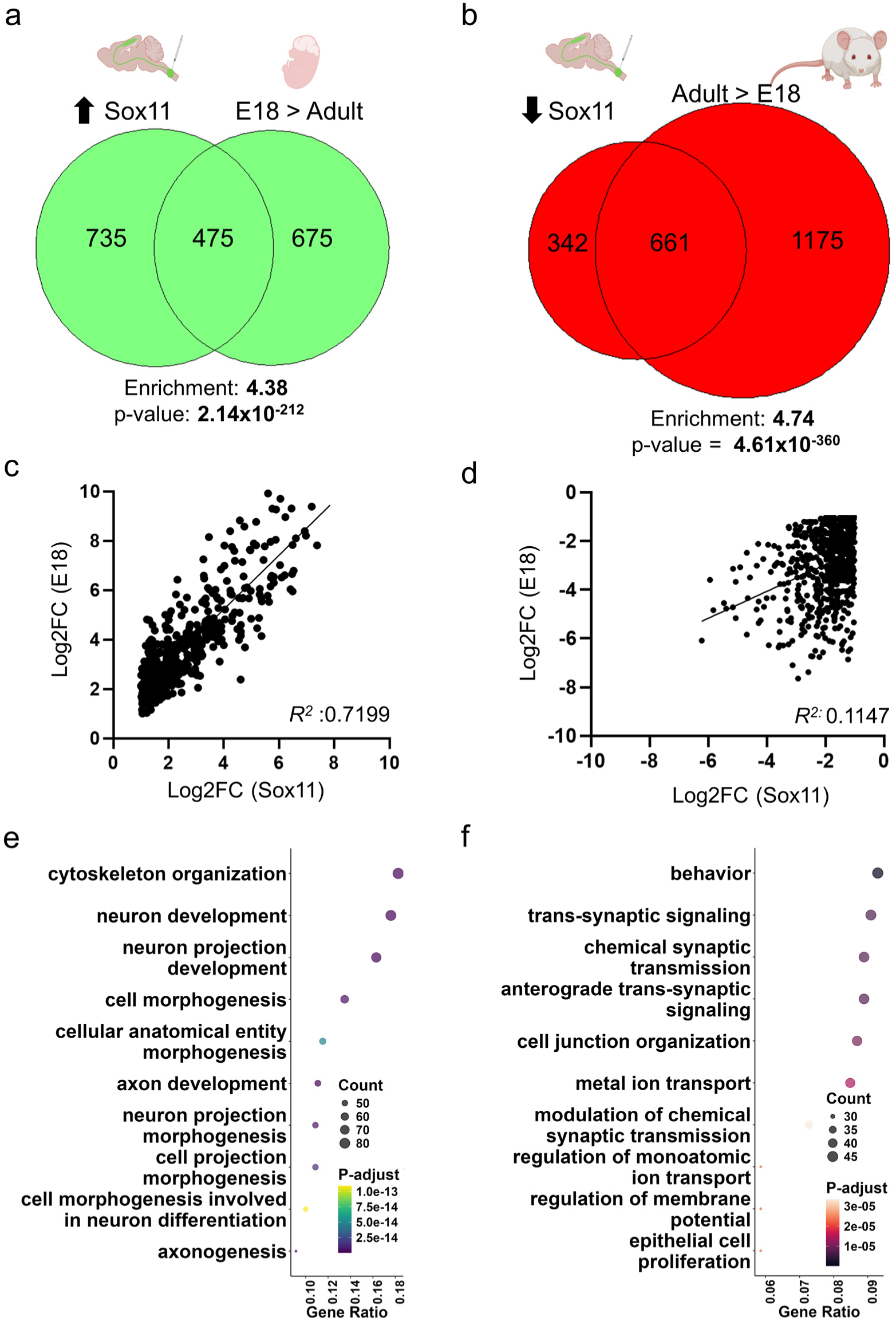
Sox11 partially restores an embryonic pattern of gene expression. (a-b) Venn diagrams showing the overlap of Sox11-upregulated (a) and downregulated (b) transcripts with embryonic and adult gene sets, respectively (p-values reflect hypergeometric tests). (c,d) Scatter plots comparing Sox11-triggered fold changes versus developmental fold changes, with linear regression analysis indicating positive correlation. (e,f) Gene ontology analysis showed biological processes that are significantly enriched in the Sox11-up / E18 overlap (e) and in the Sox11-down / Adult overlap (f) (Log2FC, ±1 p-value indicates Benjamini-Hochberg test).

## DISCUSSION

We have provided new data that clarify the transcriptional underpinnings of a key step in neuronal maturation marked by an apparent tradeoff between axon growth and synaptic activity. Remarkably, we find that forced expression of Sox11 in fully mature neurons can substantially reverse this transcriptional shift, simultaneously elevating numerous transcripts functionally involved in axon growth and suppressing transcripts linked to synapse function and excitability. Sox11’s effects are large in scale, stable through time, and occur even when Sox11 is delivered to chronically injured neurons. Together, these results demonstrate that developmental growth programs can be re-engaged in adult neurons and point toward a regenerative strategy that bridges developmental biology and pro-regenerative transcriptional programs after injury.

The single-nucleus approach used here provides updated information regarding changes in transcription that accompany corticospinal tract (CST) maturation. A prior study used microarray technology to compare murine cortical neurons between ages embryonic day 18 (E18) and postnatal day 14 (P14), yielding a widely cited set of developmental DEGs that has continued to serve as a primary point of comparison for regenerative studies [6,37,38]. Key methodological differences in the current data include the use of fully mature CST neurons and the use of snRNA-seq technology, which offers a more comprehensive detection of transcripts and the ability to exclude off-target cell types. Comparison of DEGs detected by the two approaches found high statistical concordance (3.8-fold enrichment, p-1.39×10-75, hypergeometric test) and broad agreement in GO enrichment analysis. Notably, the current snRNA-seq approach detected more than 1000 additional DEGs above a two-fold threshold that were either not present in the prior microarray data, or which were not significantly altered by the prior P14 end point. Thus, the snRNA-seq approach across full maturation provides an expanded view of developmental DEGs. An important caveat, however, is that in the present study the E18 data derives from a mixed population of glutamatergic deep-layer projection neurons and not from purified CST as previously. Thus, the present data are best suited to inform understanding of cellular behaviors that are broadly shared across glutamatergic cell types at E18, including axon projection, but are likely less sensitive to features that are highly specific to CST neurons, which comprise only a subset of the selected E18 nuclei. In this regard the developmental DEG set presented here is best considered a sensitized complement, but not a replacement, for those of Arlotta and colleagues.

Our developmental comparison largely supports the hypothesis of maturational tradeoff between early transcriptional support for axonal extension and later support for synapse function [5,39–42]. GO enrichment analysis showed abundant axon-related terms in the embryonic gene set, switching to enrichment for synapse-related terms in the adult. This maturational tradeoff is further substantiated by numerous indicator transcripts. Notable examples include *Dpysl3 (*CRMP4*), Dpysl5 (*CRMP5*), Stmn2 (*SCG10*), L1cam*, *Dcx*, *Marckls1*, and *Ncam* and its associated sialtransferases *St8sia2* and *St8sia4,* all with previously established links to effective axon elongation via cytoskeletal and/or adhesive processes, and which are highly expressed at E18 and sharply decline with maturation [27–29,32,33,43–46]. Conversely, we detected developmental upregulation of transcripts with canonical synapse functions such as *Snap25, Stx1b, Vamp1, Syt1,* and *Syt2.* Also prominent in developmentally upregulated transcripts were transcripts related to maturational changes in membrane potential and excitability such as *Slc12a5* (Kcc2), *Atp1a1, Kcnj6, Scn8a (*Nav1.6), and *Knca1.* Thus, these data help clarify the transcriptional profiles that distinguish neurons in a stage of axon elongation from neurons in a stage of mature synaptic transmission.

Regarding Sox11’s effects in CST neurons, perhaps the most striking feature is its magnitude, with more than 2000 transcripts significantly increased or decreased more than 2-fold. This response substantially exceeds a previous report of Sox11’s effects in injured RGCs [24]. Methodological differences, especially the snRNA-seq approach here versus bulk sequencing previously, could contribute to this difference. The key differentiator, however, likely involves the pre-delivery baseline. In the case of RGCs, Sox11 was delivered to cells undergoing a profound injury response, including strong upregulation of endogenous Sox11 even prior to experimental intervention [21,47]. In contrast, here we delivered Sox11 to neurons with largely unperturbed transcription and very low baseline Sox11 expression [26]. Therefore, the RGC outcome likely reflects only the additive component of exogenous Sox11 atop an elevated baseline, while the current data reveal Sox11’s full effect as it shifts the cell from initial transcriptional homeostasis.

The large effect in otherwise unperturbed neurons may be explained in part by Sox11’s established pioneer activity, the ability to bind nucleosome-bound DNA and locally increase chromatin accessibility [48]. We also found Sox11 to drive upregulation of chromatin remodelers, notably members of the SWI/SNF complex *Smarcc1* and *Smarcc2* (**Table 4-1**). Consistent with this, these same factors were previously identified as direct Sox11 targets in a cancerous cell line [49]. Thus, chromatin remodeling, both direct and indirect, likely contributes to the broad transcriptional upregulation triggered by Sox11 overexpression. The large number of downregulated transcripts is likely an indirect effect, as Sox11 is generally considered a transcriptional activator [13,50]. As one example, the downregulation of some synapse-related transcripts could be explained by upregulation of methyltransferase *Ezh2*, which was shown previously to inhibit promoters of synaptic genes [51]. Overall, we emphasize that the DEGs reported here represent a mix of direct Sox11 targets and indirect transcriptional sequalae. We further speculate that Sox11’s strong transcriptional effects are likely accompanied by widespread epigenetic modifications; future assays of chromatin accessibility and methylation can directly test this hypothesis.

A second notable feature of the Sox11 response is its high overlap with developmental regulation. More than 1100 transcripts strongly affected by Sox11 (±2-fold change) were also strongly altered during development, representing a broad shift toward a more embryonic-like pattern of gene expression. Consistent with this, GO analysis identified similar processes in Sox11 and embryonic gene sets, including upregulation of cytoskeletal organization and neuron projection development versus downregulation of trans-synaptic signaling and transmembrane ion transport. As emphasized throughout, this interpretation of Sox11’s effects are also supported by numerous examples of transcripts that are 1) functionally linked to axon growth 2) strongly down-regulated during maturation and 3) up-regulated in Sox11-expressing neurons. These include *Dpysl3 (Crmp4), Dpysl5 (Crmp5), Stmn2 (*SCG10*), L1cam*, *Dcx*, *Marckls1*, and *Ncam* and its associated sialtransferases *St8sia2* and *St8sia4* [27–29,32,33,43–46]. Interestingly, Sox11 also triggered downregulation of adult transcripts involved in synaptic release and maturational changes in excitability, including transcripts previously linked to axon growth: *Slc12a5* (Kccn2) [31], voltage-gated calcium channels, *Cacna2d2* [8] and synaptic vesicle priming genes, *Munc13s* [5]. Thus, Sox11 possess the ability to broadly reverse developmental shifts in gene expression, including changes in numerous transcripts with established roles in axon growth.

A third outcome of interest is the conservation of Sox11’s effects across cell types. Comparison of Sox11’s effects in CST neurons to RGC and epidermal studies revealed common Sox11-responsive networks involved in cytoskeletal regulation and cell projection development. Sox11 has long been recognized as a member of a small set of transcription factors whose upregulation is nearly universal in regeneration-competent neurons across species and the CNS/PNS divide [52]. Our comparative analysis helps explain this observation, supporting the hypothesis that Sox11 may drive a conserved gene module for cytoskeletal organization during axon growth. Indeed, the overlap of Sox11-responsive transcripts with epidermal cells suggests that this conservation may extend even more broadly to impact cellular motility in non-neural cell types.

Our data also reveal several Sox11 properties of high translational relevance. We show that Sox11 triggers widespread transcriptional changes within two weeks, followed by almost no further change for at least the next four weeks. This observation reinforces our prior conclusion that forced re-expression of Sox11 has no effect on cell viability in CST neurons, based on the absence of markers for cell death and no decrease in CST cell number after Sox11 delivery [53]. Consistent with this, but unlike the prior report in RGCs [24], here we detected no change in cell counts or enrichment of apoptotic-associated functions at any time point sampled. Thus, rather than driving progressive gene changes or triggering cell death, Sox11 appears to move CST neurons into a transcriptional state that is dramatically shifted from baseline but relatively stable. Finally, we found Sox11 to display the same ability when delivered to CST neurons with a four week delay post-injury, positioning Sox11 as a tool to potentially prime neurons for growth even in the chronic injury state.

Importantly, however, our results also highlight a seeming discrepancy between the large effect on gene transcription and a more modest effect on axon growth. Our prior work found that forced expression of Sox11 in CST neurons produces short distance arborization but not a full recapitulation of long-distance axon elongation after spinal injury [53]. One likely explanation involves the numerous growth-inhibitory cues in the damaged spinal cord, which were not addressed in prior studies, and which could have masked Sox11-triggered gains in intrinsic growth potential [54,55]. Thus, an attractive future direction would be to combine Sox11 as a stimulus to enhance intrinsic growth potential with strategies that alleviate cell-extrinsic inhibition. In addition, despite Sox11’s strong effects on growth-relevant gene modules, especially cytoskeletal, we emphasize that axon growth ultimately depends on a broad range of interlocking cellular processes, some of which may be transcriptionally regulated by factors beyond Sox11. For example, we found E18 neurons to be highly enriched in transcripts involved in translation and ribogenesis, which may reflect the high protein demand of rapid axon construction. However, we did not find these translation and ribosomal products among the set of Sox11- responsive transcripts. In this way, a key application of the present data will be to identify growth-relevant gene modules that fall outside of Sox11 regulation, thus narrowing the search for additional gene regulators that are most likely to complement and synergize with Sox11.

In summary, the data here reveal new transcriptional signatures that compare neurons in a stage of axon elongation to a stage of fully mature synaptic function, an important instance of the wider phenomenon of tradeoff between regeneration and differentiation through development. They also provide detailed insight into Sox11’s capacity to profoundly shift neuronal transcription into a stable pattern with striking resemblance to a prior growth-competent, embryonic stage. Importantly, Sox11 displays this ability in neurons that normally show only muted response to axon damage and in chronically injured neurons. These findings highlight the centrality of Sox11 in setting the transcriptional balance between developmental stages and re-focuses attention on Sox11-based stimulation as an influential component of pro-regenerative molecular strategies.

## METHODS

### Experimental model and study participant details

Male and female (8-12 weeks old) wild-type (C57BL/6J, The Jackson Laboratory) mice were used for this study. All procedures involving animals were conducted in strict adherence to ethical guidelines provided by the Guide for the Care and Use of Laboratory Animals published by the National Institutes of Health (NIH). The Institutional Animal Care and Use Committee (IACUC) at Marquette University reviewed and approved all animal experimental protocols (approval numbers 3283, 4013). No notable sex-dependent differences were observed during analysis. Housing conditions for the mice included ad libitum access to food and water, a controlled 24-hour cycle of 12 hours of light followed by 12 hours of darkness, temperature of 22°C ± 2°C, and relative humidity between 40% and 60%.

### AAV Preparation

AAV2-Retro-CAG-H2B-mGreenLantern was produced by the University of Miami Viral Vector Core using Addgene plasmid #177332. AAV2-Retro-CAG-Sox11 was produced by the University of North Carolina Viral Vector Core using plasmid in which H2B-mGl was replaced by DNA corresponding to the open reading frame of mouse Sox11 (accession NM_009234), synthesized by Genscript. AAV2-Retro-Malat barcode constructs were produced as previously described in [26]. Briefly, randomly generated sequences were synthesized and annealed to an RNA nuclear localization sequence derived from human Malat RNA and placed downstream of a CAG promoter. In addition to the previously described barcodes BC0, BC1, and BC102, a fourth barcode designated BC3 was constructed by replacing the BC1 sequence with randomly generated nucleotides (AAACAAGGGTCACGAGTTAAG). Barcode AAV2-Retro was produced by the University of Miami Viral Vector Core. All viruses were diluted with 1X phosphate buffered saline (PBS) (Electron Microscopy Sciences, Ref. 15710) and a protective solution of 0.001 – 0.01 % P-188 (ThermoFisher 24040032) (and 5% D-Sorbitol (Sigma, S1876) immediately before injection.

### Animal Surgeries

For spinal AAV injection, adult female and male C57BL/6 mice (6-8 weeks old, 20-28 g) were anesthetized by ketamine/xylazine (10 mg/kg), the vertebral column was surgically exposed, mounted on a custom stabilization device, and laminectomy was performed at cervical level 5 (C5). AAV2-Retro particles (0.8 µl) were injected in the right C5, 0.33 mm lateral to the midline, and to depths of 0.6 and 0.9 mm, via a pulled capillary tube fitted to a Hamilton syringe and driven by a Stoelting QSI pump (catalog #53311) and guided by a micromanipulator (pumping rate: 0.06 µl /min). For spinal cord contusion injuries, adult mice were anesthetized by ketamine/ xylazine (10 mg/kg). Mice were mounted on the custom stabilization device, and a laminectomy was performed at C5. After laminectomy, all mice received a 60 kilodyne injury at the midline of C5 using an infinite horizon impactor device (Precision Systems and Instrumentation, Ref. IH-0400). Immediately following injury, mice were placed on a heating pad and received a subcutaneous injection of meloxicam (5 mg/kg) once a day for two consecutive days.

### Embryo Dissection

Timed pregnancies between female and male C57BL/6 mice produced E18 embryos. Pregnant mothers were anesthetized by exposure to isoflurane and embryos were removed and placed in ice-cold 1x PBS. The cortex of each embryo was dissected and washed in ice-cold Ca+/Mg+-free HBSS (Gibco 14175) and promptly flash frozen on dry ice in a 1.5mL Eppendorf DNA LoBind Microcentrifuge tube. The samples were then stored at -80 °C until nuclear extraction and fluorescently activated nuclei sorting (FANs, BD FACs Melody). Before FANs, the nuclei were labeled with propidium iodide (Sigma, P4864, 1:500 dilution).

### Tissue Collection for Single Nuclei Analysis

Following previously published procedures [26], two weeks after AAV injection, mice were anesthetized with isoflurane, decapitated, and brains rapidly placed in ice-cold artificial cerebrospinal fluid (ACSF) (Hearing et al., 2013) for 1 minute. On ice, brains were then sectioned in the horizontal plane at 500 microns using an Adult Mouse Brain Slice Matrix (Zivic Instruments BSMAS005). Retrogradely labelled tissue from the left and right cortex was then micro-dissected in ACSF using a stereomicroscope and fluorescence adapter (NIGHTSEA SFA-GR). The dissected tissue was flash-frozen on dry ice in a 1.5mL Eppendorf DNA LoBind Microcentrifuge tube. The samples were then stored at -80 °C until nuclear extraction.

### Nuclear Extraction

For extraction of nuclei, frozen tissue was promptly transferred from -80 °C storage to a chilled 2mL dounce (KIMBLE, Ref. DWK885300-0000) with chilled Nuclei EZ Lysis Buffer (Sigma-Aldrich N 3408). The sample was dounced 25 times with pestle A and 20 times with pestle B. The dounced sample was then transferred to a chilled 15mL conical tube and an additional 2mL of Nuclei EZ Lysis Buffer was added and gently mixed by inversion. The sample was incubated on ice for 5 minutes and centrifuged at 500G at 4 °C. The supernatant was removed, and the pellet was resuspended in 4mL of Nuclei EZ Lysis Buffer. The sample was incubated on ice and centrifuged at 500G at 4 °C for 5 minutes. Following the centrifugation, the supernatant was removed, and the pellet resuspended in 750 µl of Nuclei Suspension Buffer (NSB) (2% Bovine Serum Albumin (BSA, Sigma-Aldrich Ref. A9418), 40U/ µl RNase Inhibitor (Invitrogen Ref. AM2684), 1X PBS). For embryonic samples, which lacked AAV-derived nuclear mGl fluorescence propidium, iodide (Sigma, P4864, 1:500 dilution) was added to NSB to facilitate sorting.

### FANs and Library Preparation

After resuspension in NSB the nuclei were passed through a 20µm filter (CellTrics 040042325) and quickly moved to the FANs machine. The collection tube for the sorted nuclei was coated with 5% BSA and contained the 10X Genomics Reverse Transcription Master Mix. Single nuclei were separated from debris and doublets using sequential gating of side scatter area (SSC-A) vs forward scatter area (FSC-A), forward scatter width (FSC-W) vs forward scatter area (FSC-A), and side scatter width (SSC-W) vs side scatter height (SSC-H), followed by fluorescence gating to gather H2B-mGl positive nuclei. With run times capped at seven minutes, between 5000 and 700 nuclei were sorted directly into the Reverse Transcription Master Mix and single-nucleus libraries were prepared using Chromium Next GEM Single Cell 3’ Reagent Kits v3.1 (PN-1000269).

### Sequencing and Data Analysis

Sequencing was performed by Illumina NovaSeq X Plus at UW-Madison Biotechnology Center. Fastq files were analyzed with the CellRanger v6.1.0 pipeline on 10X Genomics Cloud Analysis with default parameters to produce a Unique Molecular Identifier for all nuclei-containing droplets. The filtered_feature_bc_matrix output file was then uploaded for analysis in Seurat 5.0 [26]. Normalization, dimensionality reduction, and cell clustering were performed in Seurat. Cell clusters were identified by performing principal component analysis (PCA) based on 30 principal components, with significant components determined by elbow plot and jackstraw test. Nuclei were clustered using the Louvain method in the FindClusters function and visualized with Uniform Manifold Approximation and Projection (UMAP). For the developmental and Sox11 injured versus uninjured comparison, the “merge” and “FindIntegrationAnchors” functions from Seurat were used to integrate datasets. To confirm deep-layer cortical projection neurons, markers Sox10/Pdgfra were used to remove oligodendrocyte precursor cells, and Pax6/Top2a was used to remove mitotically active neural stem cells. Next, Slc1a3 (EAAT) was used to remove radial glial cells, Gad2 to remove cortical inhibitory neurons, and Cux2 to remove neurons in the shallow layer. Lastly, Bcl11b was used to identify and confirm deep-layer cortical projection neurons, thereby arriving at a final population. The remaining nuclei were re-clustered and plotted via UMAP. Cluster plots in Seurat were generated using the “scCustomize” package in R (https://github.com/samuel-marsh/scCustomize). Differential gene expression analysis was conducted using the “FindMarkers” function in Seurat, applying a non-parametric Wilcoxon rank-sum test. MA-plots were created using the “ggplot2” package (https://ggplot2.tidyverse.org/).

### Gene Ontology and Network Analysis

Gene ontology analysis was conducted using the “ClusterProfiler” (https://www.ncbi.nlm.nih.gov/pmc/articles/PMC3339379/) package in R 64,65, with input genes showing a Log2 fold change value ≥1 or ≤-1 and an adjusted p-value <0.05, marking them as upregulated and downregulated, respectively. The “org.Mm.eg.db” database served as the reference for ontology searches. The analysis included Biological Process (BP), Cellular Component (CC), and Molecular Function (MF) categories, setting gene count size limits between 3 and 500, and a p-value cutoff of <0.05. The top 15 pathways were visualized using the dot plot function from the enrichR package in R, and the results were saved as a CSV file.

### Ingenuity Pathway Analysis

(IPA; QIAGEN Inc., https://www.qiagenbioinformatics.com/products/ingenuity-pathway-analysis/) was used to analyze differentially expressed genes to identify upstream regulators. A list of differentially expressed genes, including official gene symbols, log₂ fold changes, and adjusted p-values, was uploaded into IPA. Only genes meeting an adjusted p-value < 0.05 and a log₂ fold change ≥ 2 were included in the analysis. IPA maps the input gene list to its curated knowledge base of molecular interactions, biological functions, and regulatory networks. Upstream regulators were inferred based on known relationships between transcriptional regulators and their downstream targets, with activation or inhibition predicted by the calculated z-score. Pathways and regulators with an absolute z-score ≥ 2 and p-value < 0.05 were considered significantly activated.

### Fluorescent *in situ* Hybridization

Fluorescent *in situ* hybridization was performed using RNAscope Multiplex Fluorescent Detection Kit v2 (Product #323100) from ACDbio. Mice were anesthetized by C02 exposure, decapitated, brains removed, frozen in O.C.T. Compound (Tissue-Tek, Ref. 4583), and cryosectioned at 30µm. Cryosection slides were dried at room temperature for one hour before storage at -80°C. Fluorescent *in situ* hybridization was performed following the RNAscope protocol, as follows. Immediately after removal from -80°C slides were fixed in fresh 4% paraformaldehyde (PFA) for one hour at room temperature and washed with 1X PBS twice for two minutes each. The slides were then dehydrated in 50%, 70%, and twice in 100% ethanol, five minutes each. Slides were allowed to dry at room temperature, followed by addition of hydrogen peroxide for ten minutes. Slides were then washed with nuclease-free water, twice for two minutes each. Protease III was then applied for eight minutes at room temperature and washed with 1X PBS twice for two minutes each. Probes were then applied for two hours at 40°C and washed with 1X wash buffer twice for two minutes each. Probes used were Sox11 (842061), Dpysl3 (496611), and Dpysl5 (1112131-C2). The three consecutive amplification steps were then performed, washed with wash buffer in between, as instructed by the manufacturer. The corresponding probe HRP signal was then developed. All probes were detected using a 1:2000 dilution of TSA Vivid Fluorophore 650 (PN 323273). Slides were then dried overnight and were imaged with AR1+ Confocal Microscope and 60x Apochromat Oil DIC N2 objective within two weeks from completion.

### Histology and Immunohistochemistry

Mice were anesthetized by exposure to CO2, perfused with 4% PFA in 1× PBS, brains and spinal cords were removed, and post-fixed overnight in 4% PFA. Brains and spinal cords were embedded in 6% gelatin in 1× PBS (G2500, Sigma-Aldrich, St. Louis, MO). Brains were sliced in the coronal orientation and spinal cords sagittal orientation with a vibratome to yield 100 μm sections. Sections were incubated overnight with primary antibodies Sox11 (Abcam, Ref. Ab134107), GFAP (Invitrogen, Ref. 13-030-0), rinsed, and then incubated for 2 hours with appropriate Alexa Fluor-conjugated secondary antibodies (R37117, Thermo Fisher, Waltham, MA, 1:500). Slices were then rinsed, mounted, and imaged using a AR1+ Confocal Microscope and 20x Apochromat objective. 20x images were gathered in stacked z-plane.

### Quantification and statistical analysis

Statistical details of each experiment can be found in figure legends. For snRNA-Seq experiments the number of experimental replicates and animals pooled per replicate are indicated in Results and statistical difference in gene expression was determined by non-parametric Wilcoxon rank sum tests performed within Seurat.

## ACKNOWLEDGMENTS

This work was supported by grants from NINDS R21NS135769, The Bryon Riesch Paralysis Foundation, the Christopher and Dana Reeve Foundation

## AUTHOR CONTRIBUTIONS

Conceptualization, M.B. and E.B; Methodology, M.B. and E.B; Investigation, E.B, Z.W., E.O., R.B., P.P., D.O., J.R.; Formal Analysis – M.B. and E.B; Writing – Original Draft, M.B. and E.B, Writing – Review & Editing, M.B. and E.B; Funding Acquisition, M.B. ; Resources, M.B.; Visualization – M.B. and E.B. ; Supervision, M.B.

## DECLARATION OF INTERESTS

The authors declare no competing interests.

## LEAD CONTACT

Further information and requests for resources and reagents should be directed to and will be fulfilled by the lead contact, Murray Blackmore.

## DATA AVAILABILTY

- All original code has been deposited at Github and is publicly available at https://github.com/RegenerationLab/Sox11_CST_Single_Nuc
- Sequencing data and processed data files are available through NCBI GEO Accession GSE308576 (https://www.ncbi.nlm.nih.gov/geo/query/acc.cgi?acc=GSE308576)
- Any additional information required to reanalyze the data reported in this paper is available from the lead contact upon request.

## SUPPLEMENTARY INFORMATION

Document S1. Figures S1–S7

Supplementary Table 1-1. Excel file containing additional data too large to fit in a PDF, related to Figure 1.

Supplementary Table 1-2. Excel file containing additional data too large to fit in a PDF, related to Figures 1.

Supplementary Table 3-1. Excel file containing additional data too large to fit in a PDF, related to Figure 3.

Supplementary Table 4-1. Excel file containing additional data too large to fit in a PDF, related to Figure 4.

Supplementary Table 4-2. Excel file containing additional data too large to fit in a PDF, related to Figure 4.

Supplementary Table 5-1. Excel file containing additional data too large to fit in a PDF, related to Figure 5.

Supplementary Table 5-2. Excel file containing additional data too large to fit in a PDF, related to Figure 5.

Supplementary Table 5-3. Excel file containing additional data too large to fit in a PDF, related to Table 5.

Supplementary Table 6-1. Excel file containing additional data too large to fit in a PDF, related to Figure 6.

Supplementary Table 6-2. Excel file containing additional data too large to fit in a PDF, related to Figure 6.

## Notes

### Competing Interest Statement

The authors have declared no competing interest.

