## Supplementary Data for "Sox11 overexpression restores embryonic pro-growth transcription in mature corticospinal tract neurons"

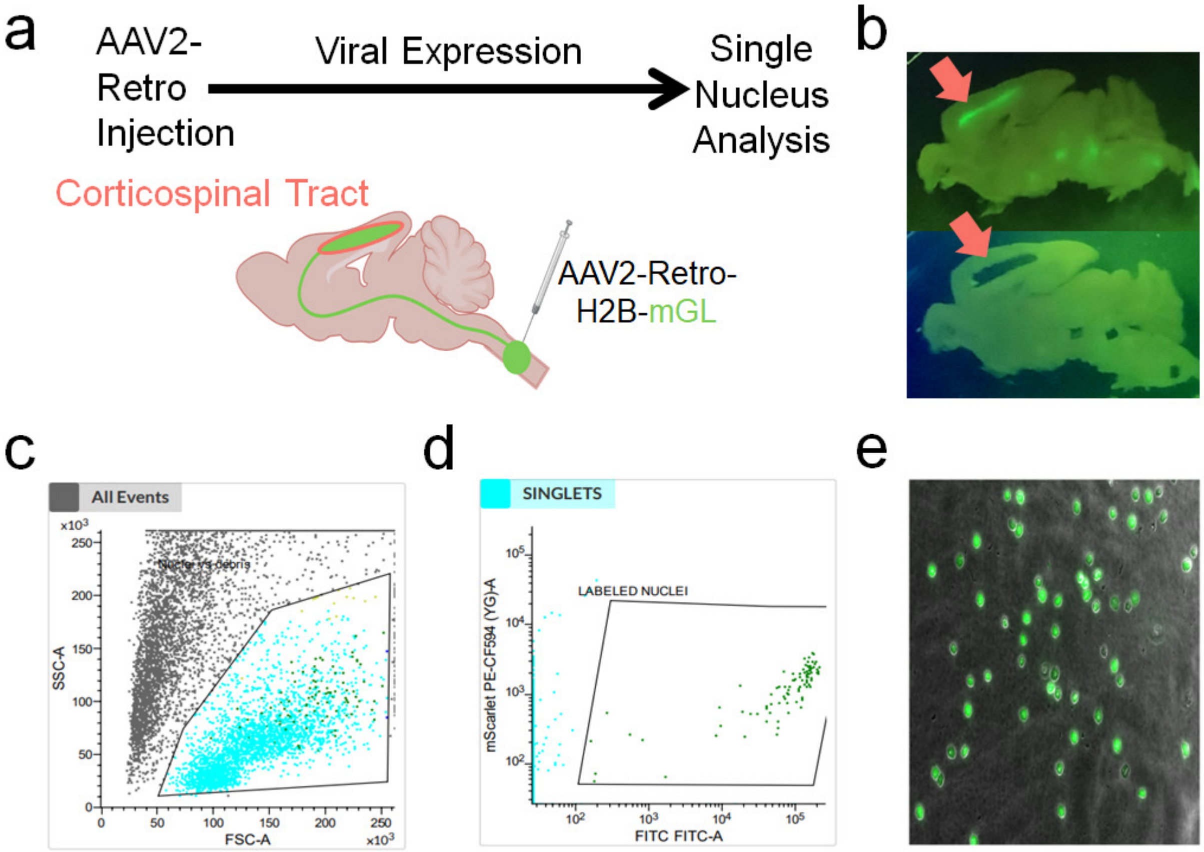


**Supplementary Figure 1-1. FANS purification of retrogradely labeled corticospinal tract nuclei.** (**a**) Experimental design in which CST cell nuclei were retrogradely labeled with nuclear-localized AAV2-Retro-H2B-mGreenlantern (mGL) at cervical level five of adult male and female mice. (**b**) Sagittal sections of brain tissue with retrograde labeled cell populations (green). Arrows indicate the region of tissue collected for CST analysis. (**c**, **d**) Scatter plots from fluorescence-activated nuclei sorting, with black lines indicating gates used to separate nuclei from cell debris (**c**) and labeled nuclei from unlabeled nuclei (**d**). (**e**) Cell nuclei after sorting with retrograde H2B-mGL signal (green), confirming high enrichment of the target population.

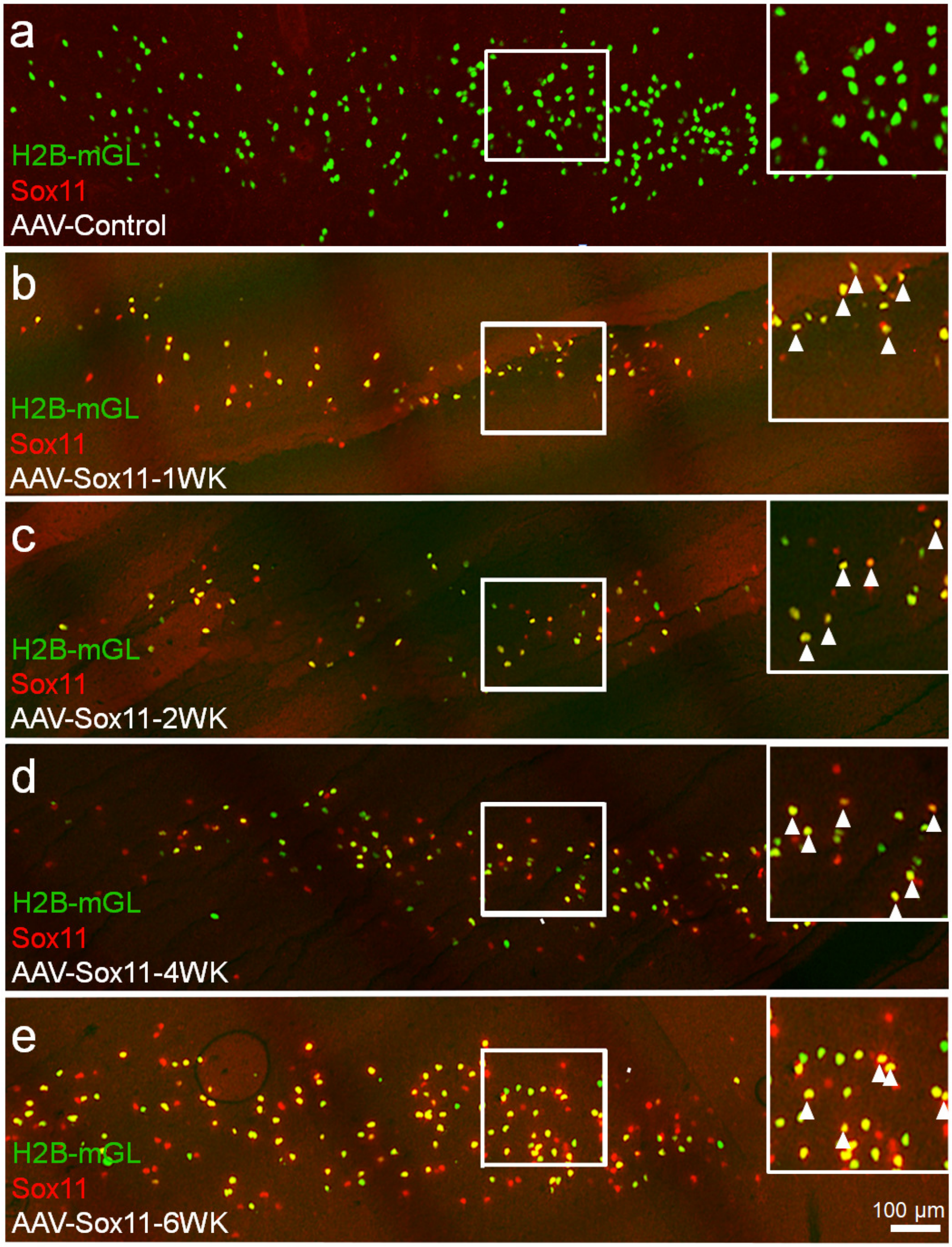


**Supplementary Figure 1-2. Sox11 is not readily expressed in the adult cortex, but retroviral delivery induces its expression.** (**a**) Shows a coronal section of cortex from an adult mouse 2 weeks after cervical spinal cord injection of AAV2-Retro-H2B-mGreenlantern and immunofluorescent detection of Sox11. CST cell nuclei (green) show no detectable signal for Sox11 (red). (**b**-**e**) Shows coronal sections of cortex one, two, four, or six, weeks after cervical injection of AAV2-Retro-H2B-mGreenlantern and AAV2-Retro-Sox11. Immunofluorescent detection of Sox11 (red) shows a strong signal in CST cell nuclei (yellow, arrows).

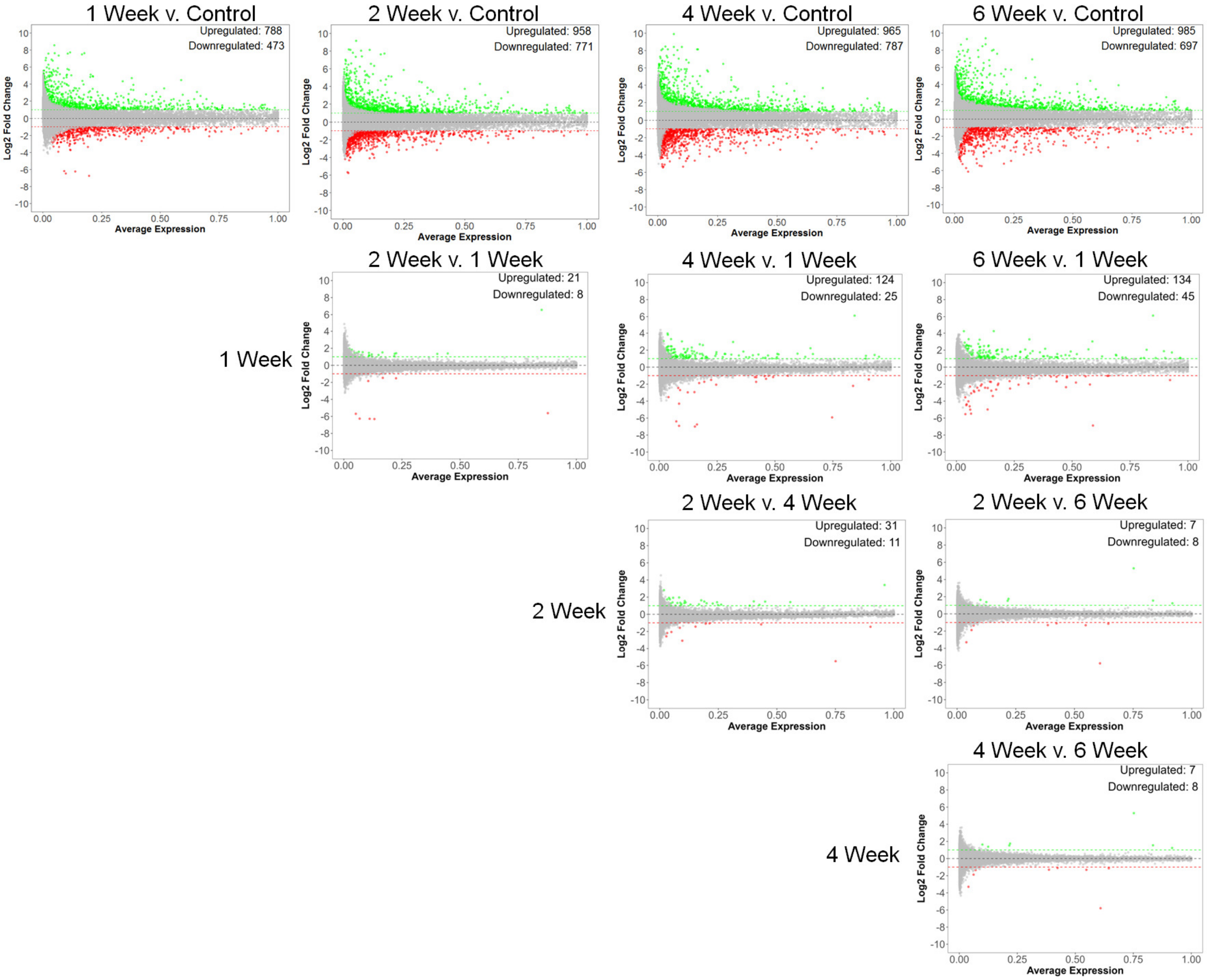


**Supplementary Figure 1-3. Sox11's effects on transcription in uninjured CST neurons are largely established and stable by two weeks post-transduction.** MA plots showing the relationship between transcript abundance and differential gene expression for two-way comparisons between Sox11 time points, revealing some differences between one week and later time points, but minimal differences between two weeks and later time points (non-parametric Wilcoxon rank sum test, p-value < 0.05).

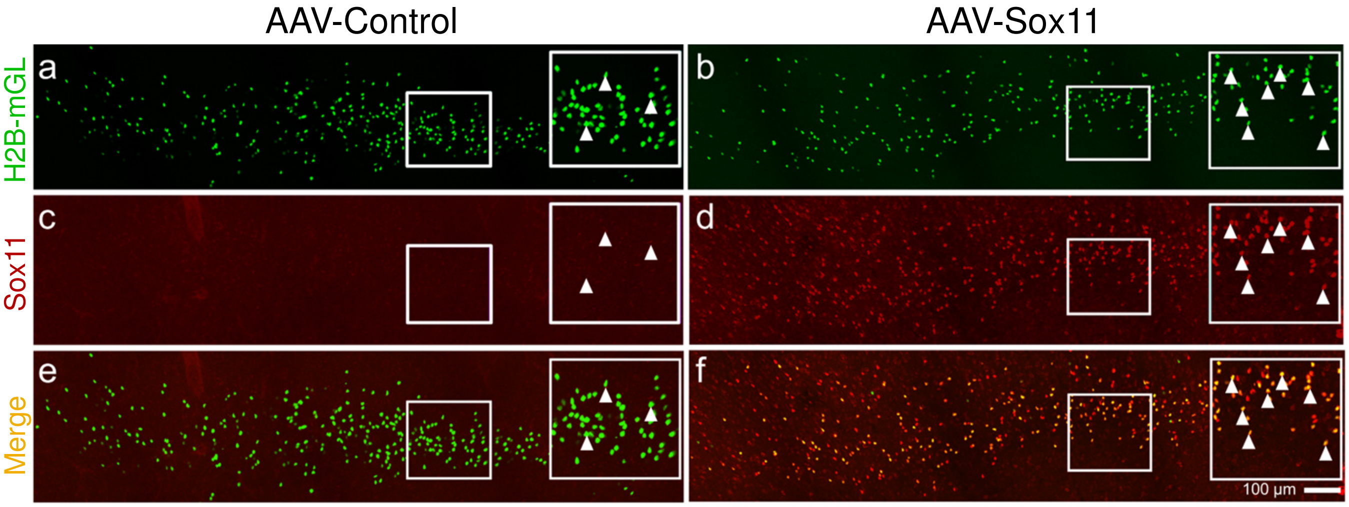


**Supplementary Figure 3-1. After a chronic cervical contusion, Sox11 is absent in the cortex, but retroviral delivery induces its expression in CST nuclei.** (**a**, **c**, and **e**) Show coronal sections of cortex from adult mice one month after a cervical contusion injury and two weeks after injection of AAV2-Retro-H2B-mGreenlantern (mGL) alone, where immunohistochemistry did not detect Sox11 (**c**, arrows) in CST nuclei (**e**, arrows) labeled with mGL (**a**). (**b**, **d**, and **f**) Shows coronal sections of cortex one month after injury and two weeks after expression of AAV2-Retro-H2B-mGreenlantern and AAV2-Retro-Sox11, where immunohistochemistry readily detects Sox11 (**d**, red, arrows) in CST nuclei (**f**, arrows) labeled with mGL.

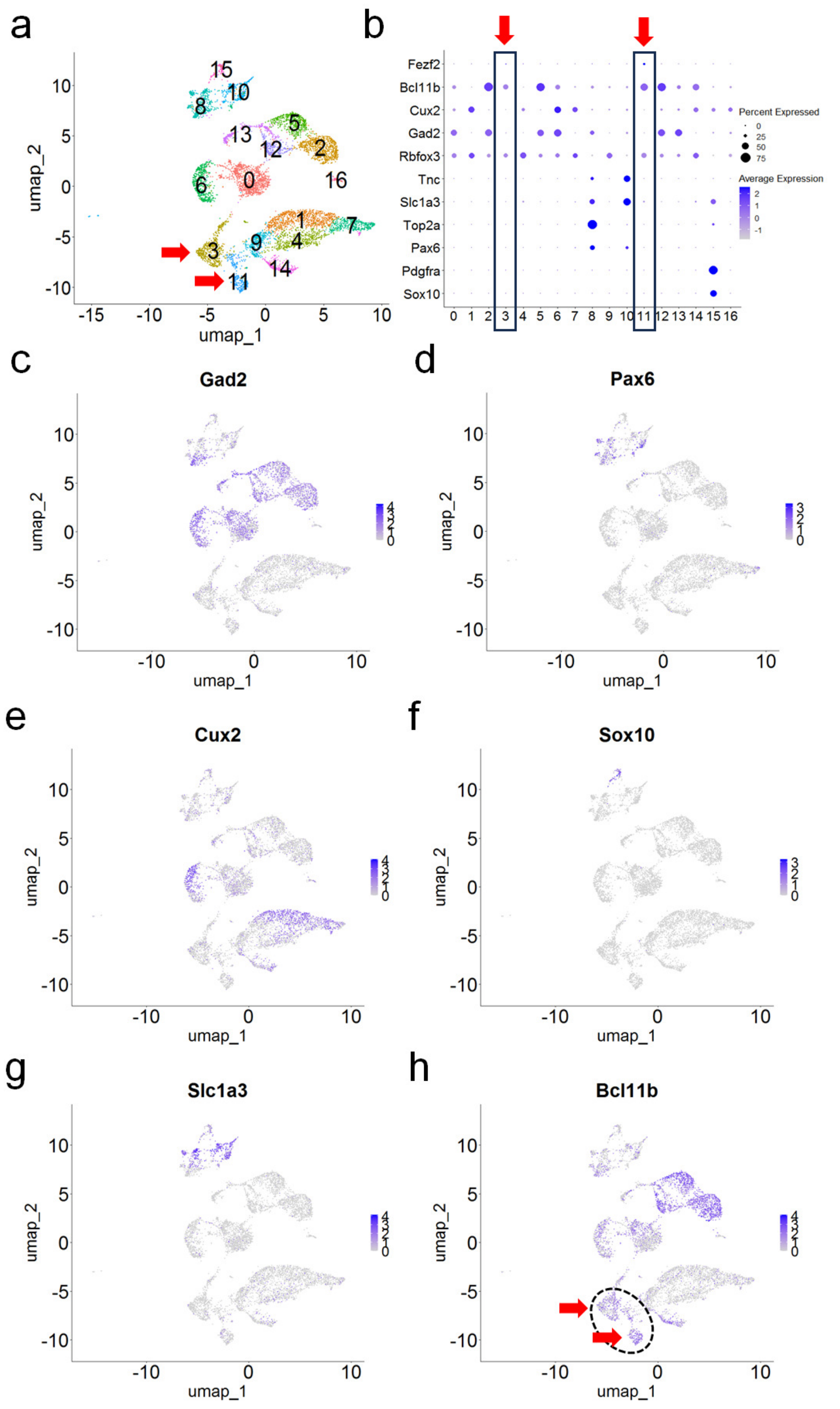


**Supplementary Figure 5-1. Embryonic deep-layer projection neurons are identified by established markers.** (**a**) UMAP clustering of 7835 embryonic cortical nuclei identifies 16 distinct groups. Arrows indicate deep-layer projection neurons. (**b**) A dotplot represents the expression of marker genes; the inset highlights clusters 3 and 11 as deep-layer cortical markers. (**c**-**h**) Feature plots show (**c**) Gad2 marks deep-layer inhibitory neurons, (**d**) Pax6 marks neural stem cells as mitotically active, (**e**) Cux2 marks shallow-layer neurons, (**f**) Sox10 marks oligodendrocyte precursor cells, (**g**) Slc1a3 (EAAT) marks radial glial cells, and (**h**) Bcl11b marks deep-layer neurons (red arrows).

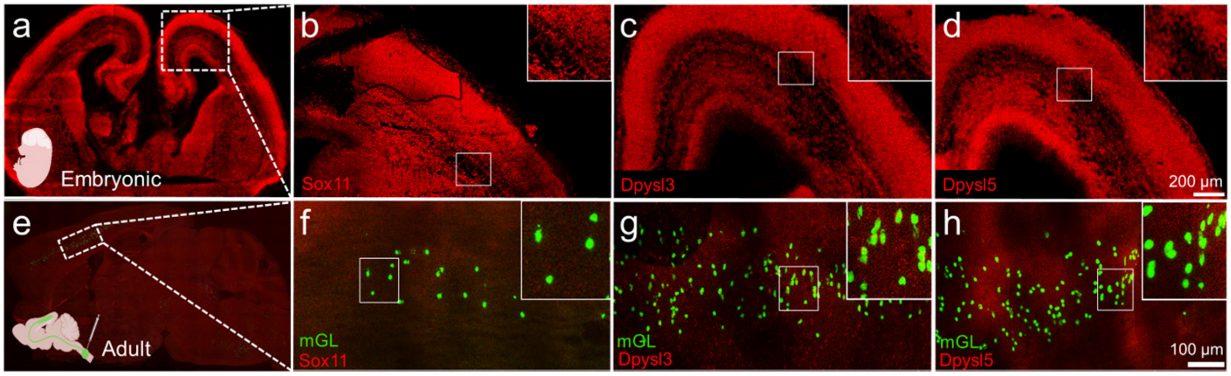


**Supplementary Figure 5-2. Fluorescence *in situ* hybridization detection confirms strong expression of Dpysl3, Dpysl5, and Sox11 in embryonic but not adult cortex.** (**a**-**d**) Show coronal sections of embryonic day 18 cortex, and *in situ* hybridization detects the presence of (**b**) Sox11, (**c**) Dpysl3, and (**d**) Dpysl5 mRNA (red). (**e**-**h**) Shows sagittal sections of cortex from adult mice, two weeks after cervical spinal injection of AAV2-Retro-H2B-mGreenlantern (mGL), and *in situ* hybridization did not detect (**f**) Sox11, (**g**) Dpysl3, and (**h**) Dpysl5 mRNA (red) in CST nuclei (green).

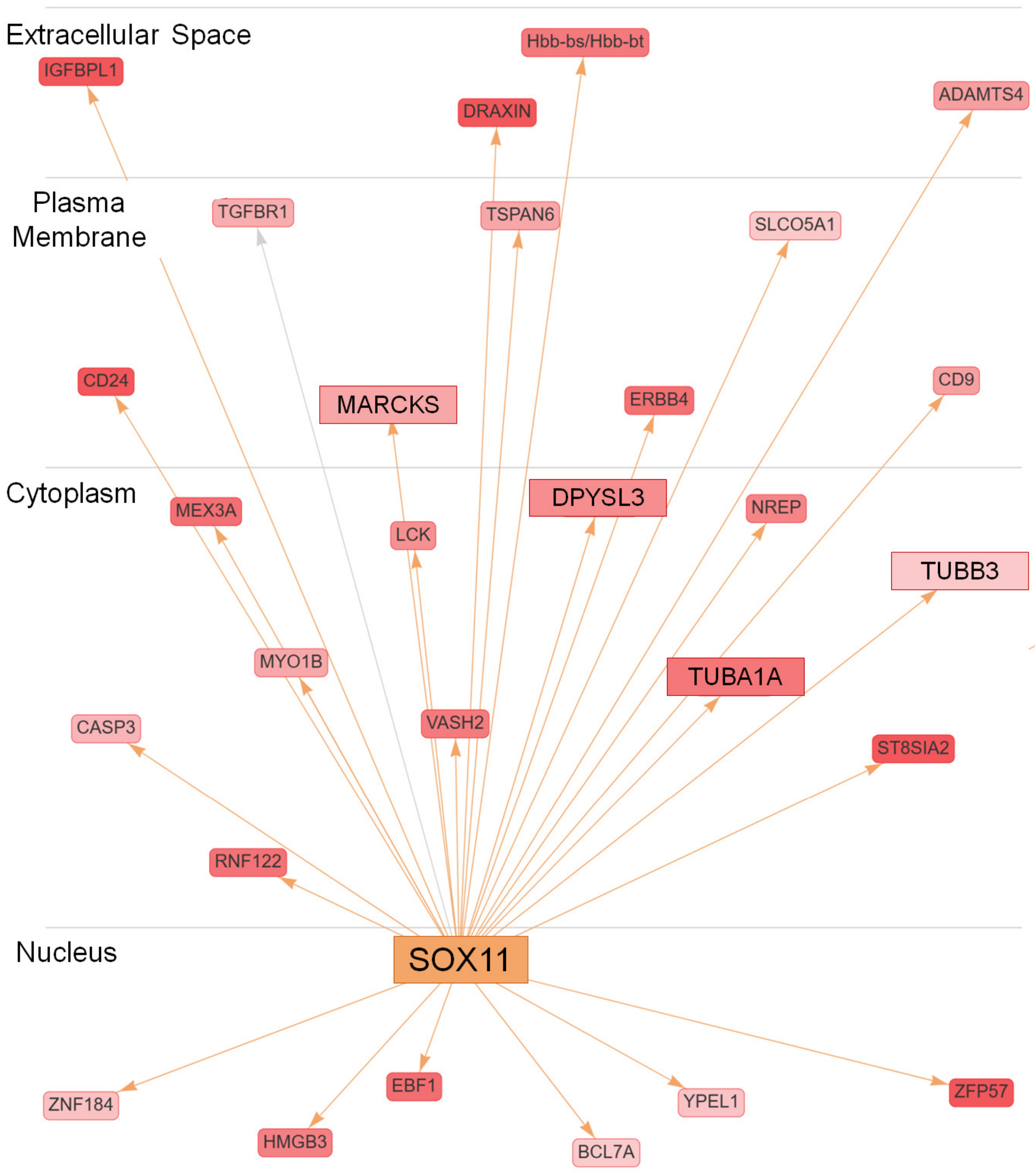


**Supplementary Figure 5-3. IPA pathway analysis predicts Sox11 as an upstream transcriptional regulator of the E18 gene set.** An interaction network from Ingenuity Pathway Analysis (IPA) ($p < 0.5$, fold change $> 2$) identifies upstream regulators of genes upregulated in embryonic nuclei. Sox11 emerged as a key nuclear regulator.
